# Foraging fruit flies mix navigational and learning-based decision-making strategies

**DOI:** 10.1101/842096

**Authors:** Sophie E. Seidenbecher, Joshua I. Sanders, Anne C. von Philipsborn, Duda Kvitsiani

## Abstract

Animals often navigate environments that are uncertain, volatile and complex, making it challenging to locate reliable food sources. Therefore, it is not surprising that many species evolved multiple, parallel and complementary foraging strategies to survive. Current research on animal behavior is largely driven by a reductionist approach and attempts to study one particular aspect of behavior in isolation. This is justified by the huge success of past and current research in understanding neural circuit mechanisms of behaviors. But focusing on only one aspect of behaviors obscures their inherent multidimensional nature. To fill this gap we aimed to identify and characterize distinct behavioral modules using a simple reward foraging assay. For this we developed a single-animal, trial-based probabilistic foraging task, where freely walking fruit flies experience optogenetic sugar-receptor neuron stimulation. By carefully analyzing the walking trajectories of flies, we were able to dissect the animals foraging decisions into multiple underlying systems. We show that flies perform local searches, cue-based navigation and learn task relevant contingencies. Using probabilistic reward delivery allowed us to bid several competing reinforcement learning (RL) models against each other. We discover that flies accumulate chosen option values, forget unchosen option values and seek novelty. We further show that distinct behavioral modules -learning and navigation-based systems-cooperate, suggesting that reinforcement learning in flies operates on dimensionality reduced representations. We therefore argue that animals will apply combinations of multiple behavioral strategies to generate foraging decisions.

## Introduction

Modular organization of biological systems provides species with flexibility to independently evolve distinct biological functions [1]. Hierarchical organization on the other hand enables coordination of multiple functions to serve common goals. Seven decades ago Nikolaas Tinbergen has recognized that animal behaviors show ample evidence of modularity and hierarchy [2]. Both experimental and theoretical work have addressed why and under what conditions distinct behavioral modules or strategies may emerge.

Spatial and temporal variations, changes in both mean and variance of quality and quantity of food patches, pose a serious challenge to all foraging animals to optimize decisions. Successful strategies must therefore be shaped to accommodate environmental uncertainty and volatility. A large body of evidence suggests that animals and humans are able to track such changes in the environment [3, 4]. In theory, animals could optimize food gathering performance by using trial and error and incorporate simple forms of reinforcement learning (RL) [5] to deal with variability in their habitats. Indeed, the RL framework has been successfully used to explain animal behavior in many learning paradigms [6]. The utility of the RL framework lies in its ability to extract decision variables that are not directly observable to the experimenter. Furthermore, using model comparison one can select the best predictive and generative RL model and see what behavioral strategies are used by animals. For example, according to standard Rescorla-Wagner RL models [6] the unchosen option values are “frozen” and updated only after the animal samples that option. Alternative RL models assume that unchosen option values decay (the animal forgets) and are updated only when the animal chooses that option. Theoretical and experimental evidence suggest that flies use the latter strategy to learn new associations [7, 8]. However, direct predictive and generative tests that would bid several RL models against each other are missing.

Besides environmental variability in their natural habitats animal are exposed to novel stimuli or new behavioral contingencies (i.e. old actions that lead to new, unexpected outcomes). According to theoretical work [9] foraging animals should explore novelty since this will lead to faster learning of behavioral contingencies. Electrophysiological recordings [10] as well as behavioral studies in mammals [11] have shown that novelty itself is rewarding. These observations can be explained by the class of RL models that explicitly incorporate novelty bonuses in the value update process [12]. Although fruit flies display behavioral and electrophysiological signatures of novelty [13], it remains to be seen whether novelty elicits behaviorally rewarding effects in flies.

Simple forms of RL-based strategies are very effective when the space of potential actions is small, but they become energetically very costly when this space grows, like in natural foraging scenarios. Therefore, alternative to trial-and-error learning animals can save time and effort by using short-cuts derived from already estimated or learned schemas [14] and spatial representations [15].

One estimation strategy is based on forms of navigation [16, 17, 18]. If a landmark can be associated with a food source, locating it can be achieved using representation of external cues [17]. In the absence of landmarks, animals use idiothetic cues [19, 16, 18] to locate previously visited rewarding locations. Furthermore, since most food in nature is not uniformly distributed, but rather occurs in patches, another strategy to maximize energy intake is to perform local searches around recently discovered food items. Indeed, local searches emerge in insect navigation when animals encounter natural [20] or fictive food sources [18, 21, 22].

Thus, navigation-based and learning strategies may complement each other by balancing efficiency and adaptability in a foraging context, to maintain and improve an animal’s fitness. Here we examine whether multiple behavioral strategies are concurrently applied by animals and how they interact. For this we designed a single-animal, trial-based probabilistic reward foraging assay in fruit flies. By dissecting the flies’ behavior into multiple behavioral modules we observed that these animals mix learning and navigation-based systems to form foraging decisions. This suggests a mechanism by which the insect brain solves the curse of dimensionality and distal reward problem faced by simple, model-free RL systems [15]. Our results imply that even in a most reduced setup, such as our plain linear-track arena, single behavioral strategies cannot be completely isolated from the ecologically sensible mixture.

## Results

### Optogenetic stimulation of sugar receptor neurons induces place preference

We set up a single-fly, closed-loop optogenetic stimulation assay where the fly is walking in a linear track arena. The trigger and reset zones are placed at each end of the arena (Fig. 1A). Single pulses of optogenetic stimulation with a fixed probability are delivered only when the fly crosses the reset and trigger zones as described in the inset in Fig. 1A. Thus, simply staying in the trigger zone will not provide optogenetic stimulation. Similar to previous studies [26, 22, 18], we observed clear effects of the light stimulation on behavior.

**Figure 1:**
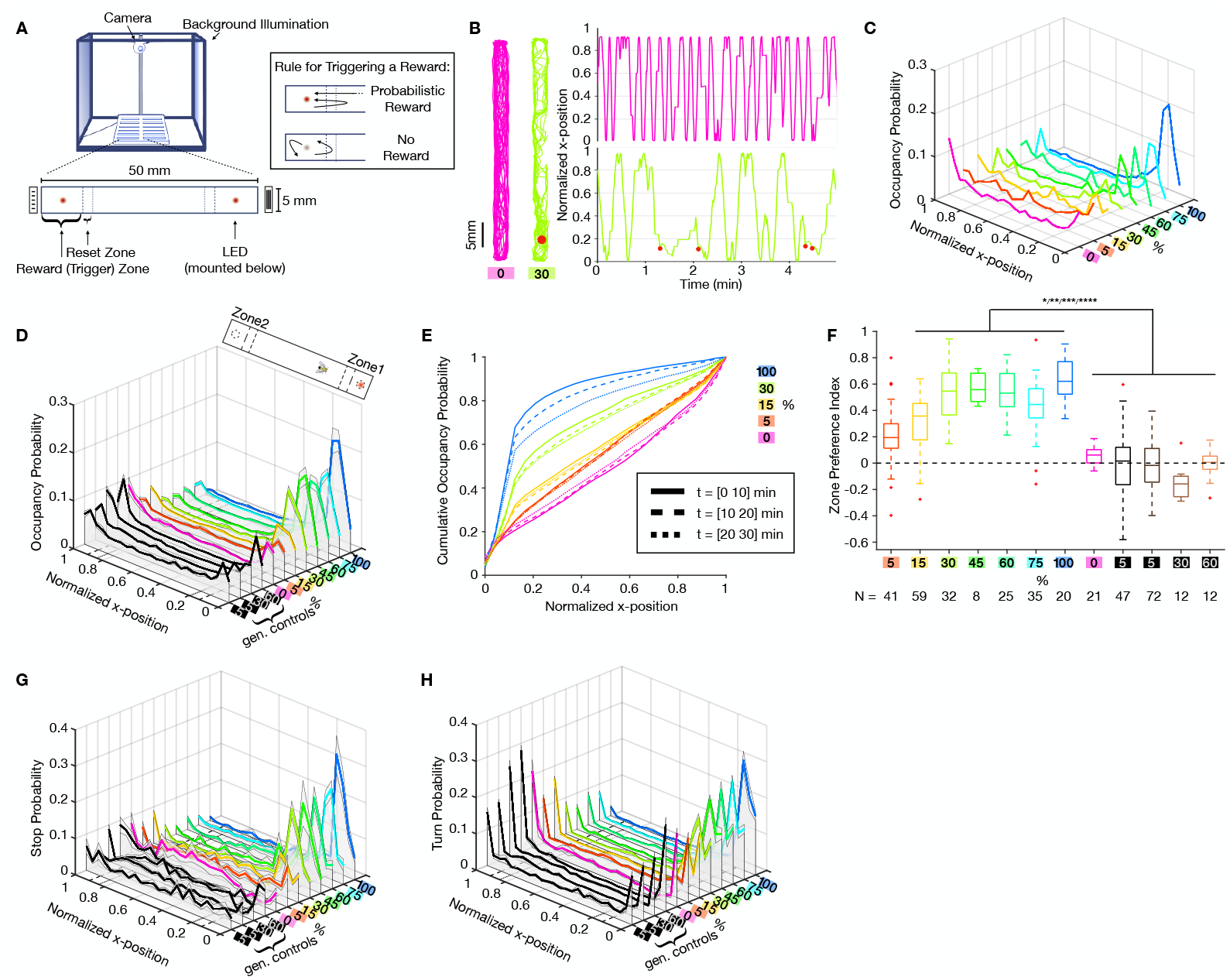
Place preference as a function of *Gr5a*-receptor-neuron stimulation. **A** Single-fly optogenetic foraging setup. A system of 12 linear track arenas is placed in a behavior box with uniform white background illumination and monitored by a webcam from above. Each arena consists of two LEDs (*λ* = 624 nm) mounted from below. Reset and trigger zones (short and long dash) are not visible to the flies. Each trigger zone is marked by black and white stripe patterns with different orientations on each side. Inset: Rule for triggering a probabilistic flash of light. Light is triggered only when the fly enters the reset and the reward zones in that order. **B** Left: Two dimensional walking traces of an unstimulated example fly (magenta) and 30% probability stimulated example fly (green). Light stimulation was delivered to only one side of the arena, here marked with red dot. Right: One dimensional walking trace over time of the same example flies. Stimulation events are marked with red dots. **C** Occupancy distribution of example individuals from 0-100% stimulation conditions. **D** Occupancy distribution of fly populations that experienced the same stimulation probabilities as in **C**. Solid lines: Mean, shaded regions around mean: ± SEM. Genetic controls don’t express Chrimson. **E** Cumulative occupancy distribution over 10 minute intervals across time. Higher stimulation probability leads to a decrease of zone preference over time. **F** Zone preference index of stimulated fly populations and genetic controls. Equal zone preference at preference index value 0 marked with black dashed line. Positive values indicate preference for zone 1 (stimulated) and negative values for zone 2. Stimulated flies have a significant zone preference of the stimulated zone over the unstimulated zone. (*: p < 0:05; ** : p < 0:01; *** : p < 0:001, Kruskal-Wallis test with multiple comparisons). **G** Stop distribution in the arena for the fly populations. Solid lines: Mean, shaded regions around mean: ± SEM. For definitions of stops and turns refer to materials and methods **H** Turning distribution for the fly populations. Solid lines: Mean, shaded regions around mean: ± SEM.

First, we looked at how kinematic variables evolved as a function of stimulation frequency. Since our setup can be seen as one-dimensional, we first focused our analysis on the walking traces along the x-axis (long axis). Optogenetic stimulation induced observable changes in the flies’ locomotion. While unstimulated, flies cover the whole space of the arena by walking back and forth, which is shown for one example fly in Fig. 1B (magenta traces). In contrast, the stimulation induced a preference for the stimulation side (Fig. 1B, green traces). By testing stimulation probabilities from 0 - 100%, we show that place preference (measured by occupancy probability of x positions) positively correlates with the stimulation frequency, both on the level of the individual fly (Fig. 1C) and the population (Fig. 1D). The 5% population occupancy distribution (Fig. 1D) is very similar to unstimulated and genetic controls, showing that the low stimulation probability is not enough to induce a significant place preference. We observe a temporal decay of the place preference (Fig. 1E) with a probability dependent magnitude. This indicates a saturation or behavioral adaptation effect in response to the optical stimulation.

To quantify the flies’ preference for the stimulation side we compared the flies’ zone preference index in Fig. 1F. Preference indices were computed from the occupancy distributions for each zone (within the reset zone boundaries), using

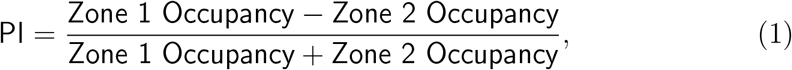

which produces preference index values between −1 and 1, meaning strong zone 2 or strong zone 1 preference, respectively, and indifference at PI= 0. There is a positive correlation of the stimulation probability and zone preference index, which is significant at stimulation probability of 15% or higher. All stimulated genetic control animals had preference indices around 0, proving that this preference effect doesn’t stem from simple attraction to the light. To test if flies had an intrinsic preference for one side of the arena that is independent of the optical stimulation, we performed double sided stimulation experiments. We observe similar levels of occupancy and place preference with these flies (Fig. S2A,B).

### Optogenetic stimulation of sugar receptor neurons triggers local search

The optical stimulation does not affect the average speed of the fly, but has an effect on speed distribution and the time duration the fly lingers around the stimulation zone (Fig. S1A, S2C,D). Thus, place preference that flies show can be due to an increase in the frequency of stop events that happens upon optical stimulation. We define stops as the speeds below a set threshold level, determined by the resolution of the camera (for precise definition of stops refer to the materials and methods section). The frequency of stops increases with stimulation probability (Fig. 1G). Increase in stop events was accompanied by an increase in turning events around the reward location (Fig. 1H). Due to the fact that the fly arena is effectively one dimensional turns are defined as velocity sing changes and indicate reversal of walking direction. The probability of turns increases upon optical stimulation on the stimulated side, while it decreases for the unstimulated side(Fig. 1H). Compared to the control population the turn frequency surpasses that at the arena walls seen in the control populations (Fig. 1H). Therefore, we concluded that local searches operationally defined as stops and turns signal local search behavior as shown in previous studies [22, 18] and suggest that the stimulation was rewarding [27].

Next we asked whether local searches simply occur as a reaction to the optogenetic stimulation (innate behavioral responses) or whether they show adaptation to the probabilistic structure of environment. We looked at the two-dimensional walking traces and computed the angular distributions of the trajectories in zone 1 (Fig. S3C). While they were significantly different for stimulated vs non-stimulated events for each probability condition, the angular distributions on stimulated trials across probability conditions were not. The same was true for probability conditions of 5 and 15% (not shown here). Together, this suggests that local searches emerge when flies receive optogenetic stimulation and they do not show any adaptation to different probability conditions.

### Flies accumulate action value over trials

In addition to initiating local searches, the animals should also return to the stimulation area, if the stimulation was rewarding. To measure this, we split the continuous walking trajectories into discrete trials. We defined a trial to be the time between two crossings of the same reset zone and the accompanying reward zone from the same direction, see Fig. 2A. This means that within a trial, the fly will have visited a reward zone, made the choice to either return to the same zone again without reaching the other zone (return decision), or to sample the other reward zone before returning. Trials also differed in whether or not the fly was rewarded when it entered the reward zone. In this way, we created a sequence of binary events given by a probabilistic reward followed by a binary choice to return or not. Fig. 2B shows the return probability for all trials (rewarded and unrewarded) to each zone (one and two) for all tested probability conditions. In all stimulated conditions, returns to the rewarded zone were significantly increased over returns to the unrewarded zone, which was not the case for the unstimulated and genetic controls (Fig. 2B, inset).

**Figure 2:**
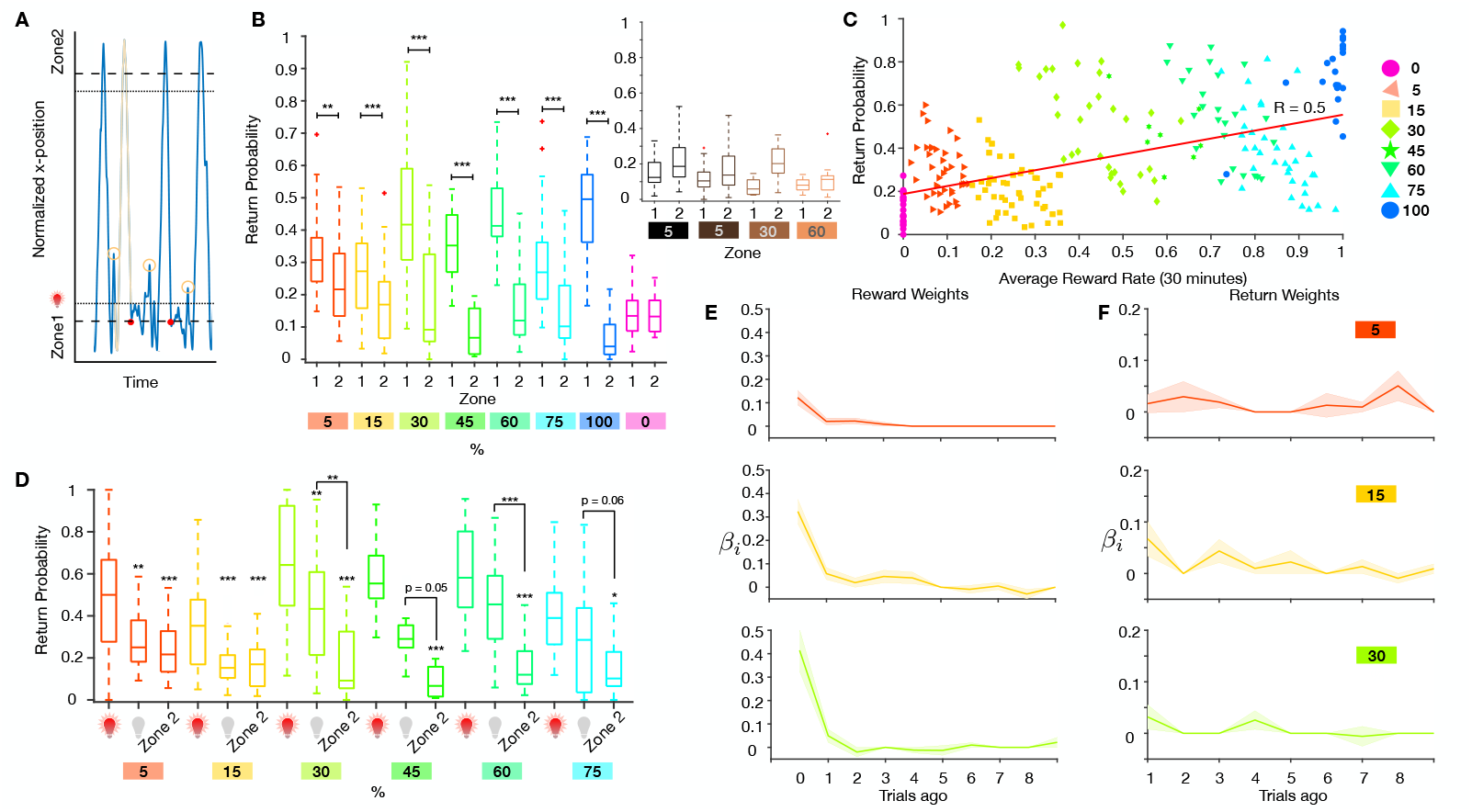
Flies return to optogenetic stimulation site. **A** Definition of a trial (highlighted trace) and a return (yellow circle). Walking trace of one example fly over time in blue. Zone boundaries marked with dashed and dotted lines. Returns are defined as trajectories that leave the reward and reset zone and return to the same reward zone, before reaching the other side’s reset zone boundary. **B** Total return behavior per probability condition to zones 1 and 2. (*: p < 0:05; ** : p < 0:01; *** : p < 0:001, Mann Whitney U test) Inset: Return behavior to both zones for genetic controls not expressing Chrimson. **C** Return probability versus average reward probability over 30 minutes. Black line: Pearson correlation, R = 0.5, p = 1e-16 (Robust Correlation package by [32]). **D** Return behavior on rewarded trials (red light bulb), unrewarded trials (grey light bulb) and always unrewarded zone 2. (*: p < 0:05; ** : p < 0:01; *** : p < 0:001, Kruskal-Wallis test with multiple comparisons). **E** Logistic regression against the reward history. Solid lines: Population averages, shaded regions: ± SEM. **F** Logistic regression of the return choices in the 5%, 15% and 30% condition against choice history for 10 trials into the past. Solid lines: Population average, shaded regions: ± SEM.

The experienced reward rate and set reward probabilities may differ due to the stochastic nature of the reward delivery. We also show a positive correlation of returns with the experienced reward rate Fig. 2C.

Since we defined returns as an additional behavioral read-out, one obvious question emerges: Are returns part of local search behavior or do they constitute a separate behavioral module? To answer this question we looked at the temporal dynamics of returns and local searches (Fig. S3A,B) and observed that while local searches are tightly locked to stimulation onset and settle to baseline within 10 − 15 seconds, returns on the other hand are mostly occurring between 15 − 25 sec. While there occasionally is some overlap, the majority of returns happens temporally separated from the local search behavior.

The trend in Fig. 2B,C suggested that flies accumulate action (return) values over rewarded trials as proposed by a previous study [22]. This effect of rewards on choices to return to the rewarded location was also seen on the strength of correlations between rewards and choices (Fig. S4B) for different probability conditions. However, our optical stimulation protocol can sensitize or desensitize directly activated neurons to subsequent stimulations. This can hinder behavioral interpretations of such manipulation experiments. Indeed, we saw strong behavioral evidence of desensitization over time in high probability stimulation sessions. The desensitization effects were most pronounced in the 100% stimulation case (Fig. S4A) and were absent at lower probabilities both for single (Fig. S4A) and double sided stimulation (Fig. S2G). This makes it look as if animals show diminishing action values. The probabilistic reward delivery allowed us to avoid this problem and analyze how returns changed as a function of reward probability, on trials where animals had not been subjected to optical stimulation. Our data shows that as reward probability and reward rate increased, returns scaled up to the rewarded location on non-rewarded trials compared to the non-rewarded locations (Fig. 2D). Thus, there is evidence that flies are accumulating an internal value of actions as a function of reward rates.

To look more directly into value accumulation we used logistic regression analysis to see how past rewards contributed to current choices and computed the reward effects (reward kernel) on return choices [28]. We show that immediate rewards had the strongest effect on current choices while rewards further back in the trial history had smaller contributions (Fig. 2E). The same effects were seen with two-sided optical stimulation (Fig. S2E, left and middle panels and Fig. S2F). Based on our analysis we concluded that flies mostly rely on current rewards to make choices, but also incorporate rewards into their choices that happen further back in trials. The simulation of fly responses to only immediate rewards generates very steep reward kernels (Fig. S4C) unlike the ones we see in the animal data. This is consistent with the idea that flies accumulate reward value over trials.

In some of the reward foraging studies using a probabilistic reinforcement structure [28, 29] not only rewards, but also past choices contribute to the animals’ current choices. This is sometimes termed decision inertia [30]. To test if flies also exhibited decision inertia we regressed current choices on past choices. Our analysis failed to detect any effects of past returns on current returns (Fig. 2F, Fig. S2E, right panel).

### Reinforcement learning models that use forgetting and learning rates capture fly behavior

Regression analysis of rewards and returns revealed that immediate rewards had strongest effects on choices. However this analysis did not distinguish how flies update the value of chosen unrewarded vs unchosen option trials (refer to the methods section for details). To see how unchosen option values are updated, we modeled the choice behavior within a reinforcement learning framework by comparing three RL models that use different update rules for unchosen options. One model freezes the value, one forgets the value with the same parameter as the learning rate *α* and the third forgets the value with a separate forgetting parameter *α*_*F*_. Examples of the evolution of the value over trials for all three models are depicted in Fig. 3A. To account for the fact that the flies have a baseline return probability below 50%, we included a bias parameter. Model selection using the Akaike informatin criterion (AIC) score [53], a measure that describes how well a model fits the data by accounting for the number of parameters (Fig. S5A), as well as predictive (Fig. S5C,D) and generative tests (Fig. S5E,F) slightly favored the second model which we termed forgetting-Q model, or FQ. The best-fit parameter values show a high variability across flies and similar mean values across experimental conditions (Fig. S5B). Predict0i.8ve testing of the best-fit FQ model on the data yielded rather poor overall accuracy (F0.6 ig. S5C,D). However, *F*_1_ accuracy reached 80% when the model was fit on data that had roughly equal, 0.2 or higher, numbers of returned to not-returned trials. Neverthe0less, under generative 5 15 30 45 60 testing the model was able to produce similar return probabilities as the%flies (Fig. 3B) and reproduced return run lengths (Fig. 3C). We defined return run lengths as the number of consecutive returns to the same side. The same analysis performed on flies with two-sided stimulation also favored the FQ model (Fig. S2H) that showed good predictive (Fig. S2J) and generative performance (Fig. S2K). However, we observed a much smaller spread of the FQ parameter values (Fig. S2I). We think that this discrepancy stems from the data limitation for one-sided stimulation trials.

**Figure 3:**
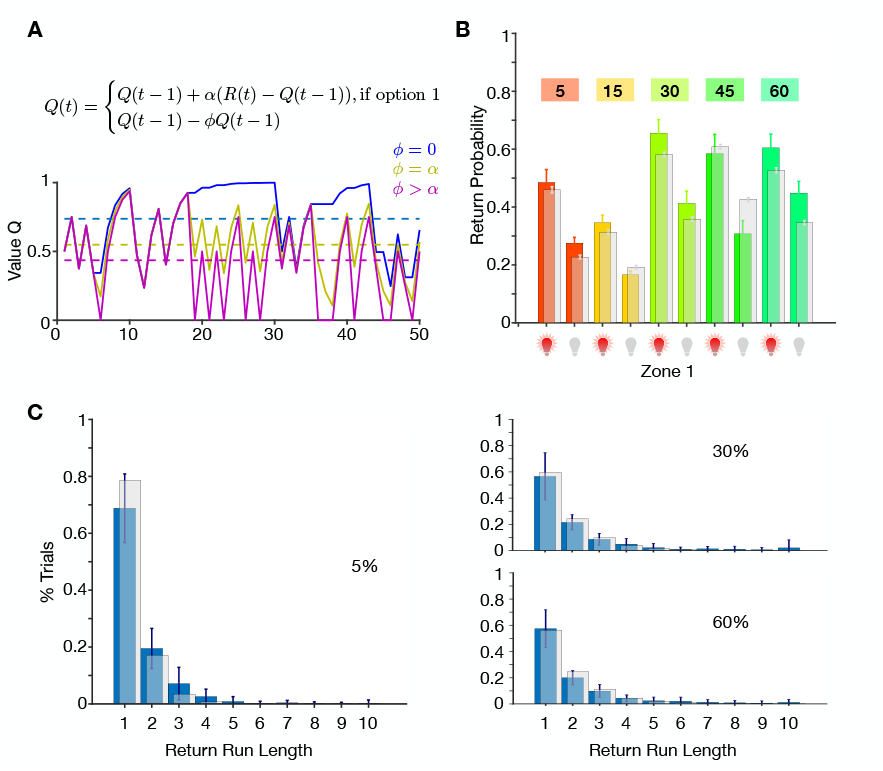
Reinforcement learning captures the probability dependent returns upon rewards. **A** Top: Update rule for the value. The three models differ in the choice of *ϕ*. Bottom: Examples of value evolution over trials for each choice of *ϕ*. RW model: blue, FQ model: yellow, FQ_*αF*_ model. **B** Generative test of the FQ model. Return probability separated by stimulated (red light bulb) and unstimulated (grey light bulb) trials to the rewarded zone per probability condition for the data (colored) and the model (grey). Bars: Population mean ± SEM. **C** Return choice run length histogram for population 5, 30, 60% fly data (blue) and the model (100 simulations, grey). Bars: Mean ± SEM. Run lengths are defined as consecutive returns to the same side.

### Fruit flies rely on cue-guided navigation in addition to trial- and-error learning to make foraging decisions

Central to all RL algorithms is that actions need to be executed before an associative learning process takes place. Alternatively, animals can execute novel actions guided by explicit representations of space and rewarded (or punished) locations [17]. To test if flies also made representation-guided choices we looked at the return probability on the very first rewarded trials. Note that due to the nature of our task design (Fig. 1A) return behavior is not required to deliver first rewards, as flies will experience rewards even if they walk back and forth the entire arena. Thus, the first rewarded trials naturally dissociate actions (returns) from outcomes, contrary to how it is done in classical operant training protocols [31]. We observed that flies returned above chance level on the very first rewarded trials (Fig. 4A).

**Figure 4:**
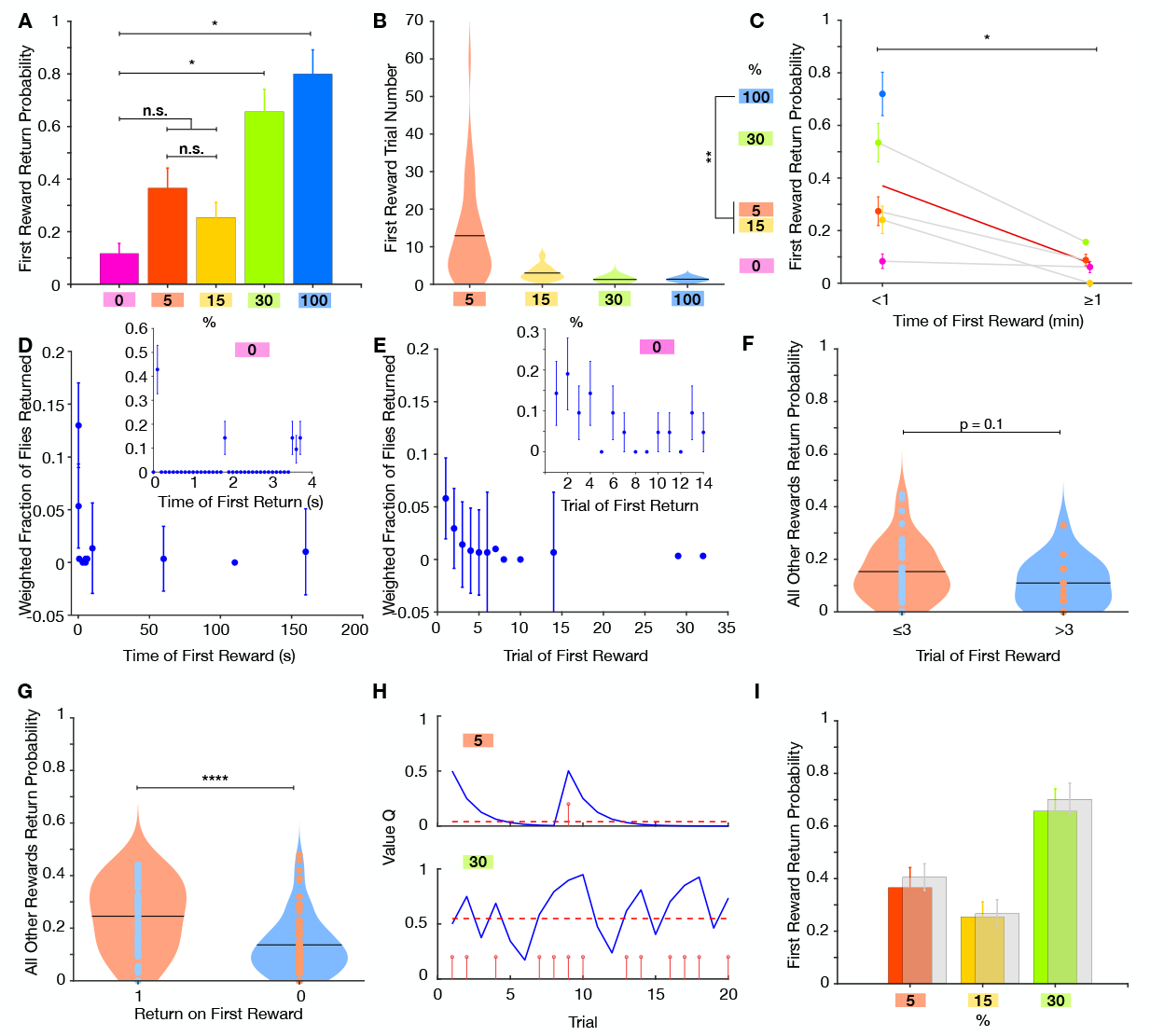
Returns on first reward. **A** Return behavior upon first reward per probability condition. For unstimulated controls (0%, magenta) the return probability was computing for the first trial. (*: p < 0.05; pairwise Fishers exact test.) **B** First rewarded trial number per probability condition. Black lines: mean. **C** Return behavior upon first reward within the first minute of recording and after the first minute. Blue circles without a connecting grey line correspond to conditions where the first reward always happened within the first minute. Red line: average return probability. Magenta circles and dark grey line: unstimulated controls. Error bars: ± SEM. (*: p < 0.05; ** : p < 0.01; *** : p < 0.001, Kruskal-Wallis test with multiple comparisons and Welch’s t-test). **D** Fraction of flies (independent of stimulation probability) that returned to a first reward within the first 200 s (time bins: 0.03 s between 0 and 1 s, 0.1 s between 1 and 9 s, 50s from 10 to 200 s and 100 s from 300 to 1000 s). Inset: Fraction of unstimulated flies that returned for the first time since the session start against time (time bins of 0.1 s). **E** Fraction of flies that returned to a first reward within in the first 30 trials. Inset: Fraction of unstimulated flies that returned for the first time against the trial index. **F** Return probability to all other rewarded trials when the first reward happened within the first 3 trials (orange violin) or after the first 3 trials (blue violin). **G** Return probability to all other rewarded trials depending on whether the fly returned upon the first reward (orange violin) or not (blue violin). (* * * : p < 0.001, Welch’s t-test) **H** Two examples for the evolution of the RL value Q (blue curve) over trials. Upper figure: 5% stimulated fly. Lower figure: 30% stimulated fly. Red stems: stimulation events. Red dashed line: average value. **I** Transparent grey bars: RL model’s prediction of first reward return rate. (Color bars as in **A**)

In our foraging assay the walls of the arena were covered with stripes that can aid the flies to navigate and locate the rewarded locations. In the studies by [16, 22] it was shown that visual or tactile cues in addition to idiothetic cues help animals to locate the rewards. This suggests that fruit flies in our assay used cue-based navigation (explicit or implicit) in addition to trial-and-error learning (simple forms of RL-based learning) to form choices.

### Novelty increases action values

We observed to our surprise the reward probability dependence of the returns on the first rewarded trials (Fig. 4A). One possible explanation could be that the flies were sensitive to the timing of the first reward in the session. To further elucidate behavioral mechanisms that drive this form of adaptation we looked at the delay of the first rewards from the start of the session using both trial (Fig. 4B,E) and time-based analysis (Fig. 4C,D). Both measures showed a similar trend of decreased return probability with increased delay. This analysis suggests that optogenetic rewards become more attractive when they are delivered in novel environment (fly arena and its edges are novel at the beginning of the behavioral session). We observed the same effect of the first rewarded trial on return probability when both sides were used to deliver optogenetic stimulation (Fig. S2D). The novelty of the arena on its own also exerted rewarding effects on flies as control flies that never experienced rewards showed above baseline level of returns that decreased to baseline (Fig. 4D,E, insets). This decay was fast as no change in returns were observed on first and later trials (Fig. 4C, magenta circles connected with grey line) on a time-scale of minutes.

Our previous analysis suggested that the action value accumulates as the number/probability of rewarded trials increases (Fig. 2D). If so, the timing of the first rewarded trial may have affected the subsequent return probability as the action value should be higher for early vs later occurring first rewards. For this we looked at the return probability on all subsequent trials when first rewards happened within the first 3 trials (this number was chosen since the return probability does not change if the fly is rewarded after the 3rd trial (Fig. 4E)) or later (Fig. 4F). We could not detect a significant difference in return probability between these groups, suggesting that except for the first few trials the behavior of the animal was not affected by the timing of the first rewards. There could be individual differences to novelty that may indicate the flies’ sensitivity to rewards in general. Therefore, we separated flies that showed return on first rewarded trial from flies that did not return on the first rewarded trial and looked at the return probability on all subsequent rewarded trials (Fig. 4G). We show that return behavior on the first rewarded trial is a good predictor of future returns and may reflect individual differences among flies.

To formally account for the observed responses of flies on the first rewarded trials we incorporated this in our RL models and assumed that option values (in our case zone 1 and zone 2 of the arena) are not set to zero initially (due to novelty), but rather start with some default positive value that over time decays (Fig. 4H). Note, that this simple model qualitatively explains novelty attraction in control flies that never experienced optical stimulation. We also tested if such RL model could correctly predict flies returns to first rewards. Our modified FQ RL model indeed generated similar return probabilities (Fig. 4I), explaining novelty mediated reinforcing effects of optical stimulation.

### Cooperation of learning and navigation-based systems

After discovering that flies apply both navigation and learning based strategies to locate the optogenetic rewards, we asked how these two systems interact with each other. Previous work [16] has shown that inbound paths of flies to their feeding sites are more straight then outbound paths, suggesting that path integration mechanisms help animals reach their feeding sites using shorter routes. Here we used a similar approach and decided to look at how flies navigated towards and away from their rewarded location as a function of accumulated value. We looked at the angular distribution of walking paths on rewarded and non-rewarded trials as a function of reward probability. The more curved a path is, the more uniform the corresponding angular distribution gets, which translates into a higher angular distribution entropy (Fig. 5A). First we noted that out-walking (walking away from the rewarded location) paths generally had slightly higher spread in angular distribution compared to in-walking paths (walking towards rewarded location) (Fig. 5B,C). This difference did not reach statistical significance. We noted, however, a consistent trend in the reduction of angular distribution of in-walking paths as a function of the reward probability (Fig. 5D) (p < 0.05 for 5 and 15% reward probability compared with the 100% reward probability). Thus, flies choose to walk more straight paths towards the rewarded locations as a function of reward value. Based on our results and the published work we speculated how learning and navigation-based systems interact at the neural level (Fig. 5E). We propose that dopamine mediated reward prediction error assigns values to spatial representations (external or internal spatial cues stored in the insect central complex [17, 33, 34] or mushroom bodies [35]). Thus, animals do not have to learn entire action sequences and instead can compare values of short-cuts to choose the better options. This effectively reduces the complexity of the action-outcome contingencies during the learning process.

**Figure 5:**
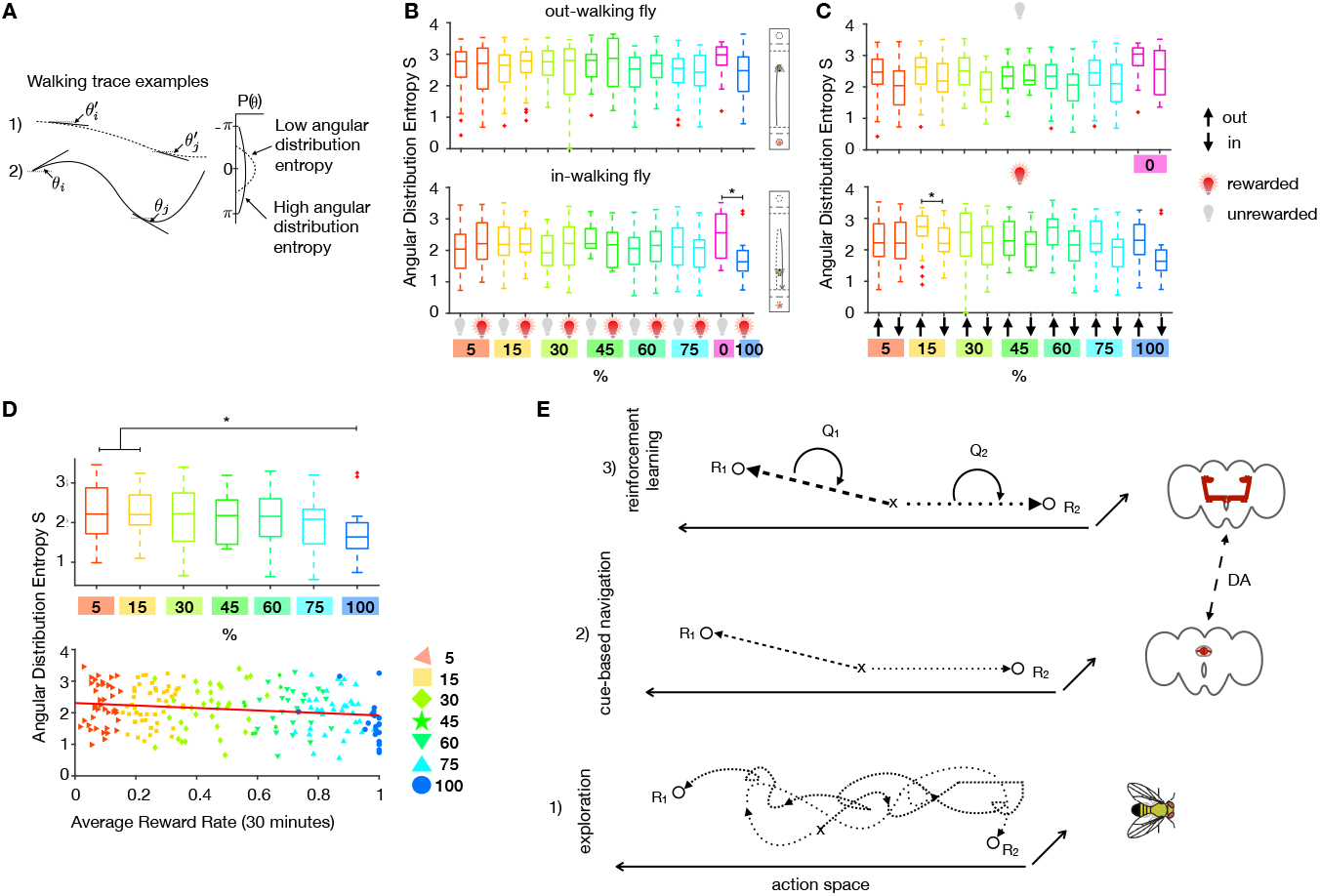
Interaction of navigation and learning systems during a foraging task. **A** Sketch of path angular distribution analysis. The ‘straighter’ path 1) is characterized by a narrow angular distribution, while the more curved path 2) has a broader angular distribution. If the distribution is narrowly peaked, it has a smaller entropy S than a broader distribution. **B** Angular distribution entropy of in- and out-walking paths, on rewarded (red light bulb) and unrewarded (grey light bulb) trials. For out-walking (away from the rewarded zone, measured from the reset zone) trajectories, there is no difference between rewarded and unrewarded trials. For in-walking trajectories (from the position of return to the reset zone), the rewarded trials have a smaller entropy, corresponding to more straight paths. This is significant for 100% compared to unstimulated controls (*: p < 0.05, Kruskal-Wallis test with multiple comparisons). **C** Same data as in **B** but sorted by unrewarded (top) and rewarded (bottom) trials. (*: p < 0.05, Kruskal-Wallis test with multiple comparisons). **D** There is a trend of more straight in-walking after a reward (indication of path integration) with increased reward probability. Top figure: only rewarded trials from **B** bottom (*: p < 0.05, Kruskal-Wallis test with multiple comparisons). Bottom figure: Same data plotted against each flies experienced average reward rate. Red line: Pearson correlation, R = −0.18, p=0.005 (Robust Correlation package by [32]). **E** Proposed schematic of the interaction of navigation and learning systems in a foraging task. 1) A foraging fly starts navigation in a new environment with the sequence of actions (dashed) that leads to reward R1. After that, the fly continues to forage on a path (dotted) experiencing another reward R2. 2) After leaving R2, the fly can make a decision to return to the R1 or R2 rewarded site via the already executed and rewarded path (dashed or dotted) or, using cue-based navigation, travel on shortcuts to the rewarded locations. 3) Combined with reinforcement learning, values are assigned to those shortcuts and updated with the collected rewards. Thus, instead of storing the entire sequence of actions, the fly needs to compare only the values Q1 and Q2 for those shortcuts, thereby reducing the complexity of the representation of their habitat. We propose a tentative biological implementation of these two processes based on previous work. We speculate that dopamine signaling assigns values to spatial representations computed in central complex (idiothetic) or mushroom bodies (external cue-based navigation systems).

## Discussion

We developed a single-fly, trial-based optogenetic reward foraging assay and discovered that foraging decisions in fruit flies can be broadly categorized into navigational and learning-based systems. Detailed analysis allowed us to discover distinct behavioral modules: local searches, cue-based navigation, novelty seeking, learning and forgetting processes. What is the biological function of these modules and how do they interact with each other in the context of foraging decisions?

As previously described for natural rewards [36] the flies in our assay initiated local searches as a function of experienced optogenetic stimulation, showing that artificial stimulation of sugar receptor neurons recapitulated the natural behavioral response in these animals. Looking at the temporal dynamics of local searches, we see that they persist for roughly 10-15 seconds after a stimulation and do not show any dependence on the frequency of the stimulation. This suggests that sweet-taste induced local search is a hard-wired behavioral module. We speculate that this behavioral module serves to anchor animals around recently discovered food items to maximize the energetic gain from that source [37].

Local searches are a useful behavior once the animal has discovered a food source. Finding a new food patch or returning to already discovered ones requires alternative foraging strategies. In insects these strategies can arise either from representation-based systems [19], using external or idiothetic cues, or learning action-outcome contingencies [38, 31]. Clear separation of these two mechanisms requires monitoring of animal behavior from the initial phase of learning to its stable performance. By controlling reward delivery with optogenetic means, we were able to track the animals’ decisions to locate rewards on the very first trial. This excluded the possibility of learning action-reward contingencies to guide animals’ choices. Our results demonstrate that fruit flies rely on representations to locate the rewarding sites, since we saw a modulation of walking paths before and after receiving optical stimulation. Representation-guided foraging decisions have a clear advantage over simple forms of learning. In stable environments representations allow animals to take novel paths, make short-cuts that save energy and minimize exposure to their predators. It is worth noting that we did not manipulate external stimuli to disambiguate contributions of external or internal cues and therefore this remains an open question that future studies can address.

However, in changing habitats animals need to learn new contingencies and, therefore, foraging decisions need to incorporate learning processes. We show that a simple Q-learning model [5] that uses both learning and forgetting rates for chosen and unchosen options, respectively, can reproduce the return behavior of the flies. Furthermore, we reveal the reinforcing effects of novelty and incorporate it into our RL framework. The new finding [13] that dopamine neurons report novelty in flies and the fact that at least some of these neurons mediate rewarding effect in flies [39] is consistent with our findings that novelty itself is rewarding. Indeed reinforcing effects of novelty might be a common driving force for exploration across species as the same phenomenon was reported in rodents [40] and monkeys [41].

What do we gain by dissecting behaviors into multiple modules? First, we can see if and how these modules interact at the level of behavior and reveal their hierarchical structure. Second, it can inform us what type of connections exist at the neural level. For example, if behavioral modules interact on the same level with antagonistic or synergistic effects, this suggests mutual inhibitory or excitatory connections exists between brain areas that control behavioral expression of those modules. Alternatively, if there are hierarchical dependencies this would suggest neuromodulatory influences.

According to our data, the RL system operates as a layer on top of the navigation-based system and a neurobiological substrate in flies exists to suggest that the RL system exerts neuromodulatory effects on the navigation-based system. The central complex monitors angular orientation in fruit flies [42]. Both neural recordings [33, 43, 34] and behavioral manipulations [44] suggest that flies use a representation-based navigation system. The neurons in the central complex (CX) express dopamine receptors [45] and these neurons control angular motion in flies [46]. Therefore, the navigation-based representations in CX can be updated (modulated) via a dopamine-mediated reward prediction error that can implement model-based learning.

We note clear similarities of hierarchical and modular structure of behavioral functions in flies and what has been first theorized and then experimentally tested in humans. Some of the computational models in RL field explicitly distinguish model-free and model-based learning systems [15]. The model-based RL framework [47] suggests that the task structure and/or spatial representations are updated by reward prediction error. This framework has been successfully used to explain hippocampus dependent changes in choice strategy when human subjects were asked to make decisions based on learned representations [48]. However, whether animals also rely on model-based learning is an open question. Some of the studies in rodents are in favor of such systems [49], yet extensive training protocols needed to achieve stable performance in animals raises doubts whether model-based learning is replaced by model-free learning system. Showing that navigation and learning systems cooperate in fruit flies, our work is consistent with the idea these animals deploy model-based learning to reduce the high dimensionality of the action space and achieve both efficiency and adaptability.

Finally, we would like to caution against reductionism in behavioral neuroscience. Recently it has been argued that neuroscience relies too much on a reductionist bias [50] in understanding the link between the brain and behavior. Here we would like to argue that behaviors themselves are subject to a reductionist bias by the desire of the experimenter to place it within a single conceptual framework. Our approach tried to break this trend and look at behaviors as composed of multiple modules, be it reinforcement learning, cue-based navigation or innate and hard-wired foraging strategies. We would like to argue that even in highly constrained environments set to focus on a particular aspect of behavior, their inner multidimensional nature should not be ignored but rather examined in detail [51]. Just to illustrate this point a recent study by Stern et al [22] argued that a spatial task in flies is solved by trial-and-error learning while Corfas et al [18] suggested that, in a very similar behavioral paradigm, animals locate rewarding sites by using a path integration mechanism. We believe that both strategies are indeed concurrently used in flies.

## Materials and Methods

Single *Drosophila* melanogaster males were starved and placed in a linear track arena, see Fig. 1A, which they were free to explore. The trehalose sugar-receptor neurons *Gr5a* [23] were chosen to express the light-activated ion channel Channelrhodopsin Chrimson [24], by means of the *LexA-LexAop* system. For details on fly strains and rearing, see the supplementary methods section.

The optogenetic fly foraging setup consists of a 3d-printed platform with 12 linear arenas of 5 by 50mm, each for a single fly, similar to Ref. [25]. The arenas are each separated by black barriers to reduce visual contact to neighboring arenas. Red light LEDs (λ = 624(631) nm, Vishay VLCS5830) are mounted from below to illuminate the respective region through a thin layer of plastic. The setup is surrounded surrounded on three sides by acrylic panels (EndLighten, Acrylite), each lit by a strip of white LEDs mounted along the end to provide white uniform background illumination and a water reservoir for humidity. The setup is monitored from above with a webcam (LifeCam Studio, Microsoft), fittedwith a short-pass filter (FESH0600 Thorlabs) to block red light from the stimulating LED. Centroid fly-tracking and stimulation are controlled in an on-line fashion by custom written MATLAB (Mathworks) scripts. In the camera view at the ends of each arena additional ROIs are defined to separate ‘reward’ and ‘reset’ zones. Using two zones allowed us to avoid self-stimulation when the fly simply stayed in the rewarded location. The reward and reset zones extend 6 mm and 3 mm, respectively, and zones of the same type are of the same size. Probabilistic rewards are triggered when the fly crosses the reset zone and enters the reward zone, in that order. Refer to the inset of Fig. 1A for a depiction of the trigger rule. The stimulation duration was 0.05 seconds.

### Fly Strains and Rearing

Flies were housed under a 12 h:12 h light:dark cycle at 25◦ C and 60–70% humidity on cornmeal, oatmeal, yeast and sucrose food. For all experiments 3-6 day old males were used, which were starved for 10-12 h prior to testing, while supplying water via a wet cloth. Flies were then transferred to the arena using an aspirator and left in the arena for 2-10 hours. The following strains were employed: *Gr5a-LexA* (gift from Kristin Scott [52]), *LexAop-Chrimson* ([24], w1118; P{13XLexAop2-IVS-CsChrimson.mVenus} attP40, Bloomington 55138), *Canton S* (from A.v.Philipsborn). The flies expressing *Chrimson* were fed all-trans retinal (ATR, Sigma Aldrich, CAS Number: 116-31-4) for 2-3 days before the starvation period. ATR food was prepared by mixing normal food with ATR to reach a 400 *µ*M solution and then covered with aluminum foil to avoid degradation. Flies fed on ATR food were kept in the dark under aluminum foil cover.

### Experimental Conditions

*Chrimson > Gr5a* flies were tested in eight different single-sided stimulation conditions; with 0, 5, 15, 30, 45, 60, 75 and 100% stimulation probability. In a second series of experiments, flies were tested under double-sided stimulation conditions, with 5-5% and 15-15% stimulation probability.

### Post-processing of Walking Data

Walking traces were cleaned of missing data points and jumps in the centroid contrast tracking and filtered with a butterworth filter using a cutoff frequency determined from camera jitter. Next, a trial structure was defined and data from flies with less than 50 trials was excluded from further analysis.

### Definition of Observables

Stops were defined by speeds below a value of |*v*| ≤ 0.01mm/s, which is governed by the resolution of our tracking system and corresponds to a movement of less than one pixel between two timestamps. Turns were defined by velocity sign changes since our setup is effectively one-dimensional.

### Logistic Regression

Regression analysis was performed on return choices against their reward history for individual flies and fly populations by averaging over individuals from the same experimental condition. Due to the binary output variable we used logistic regression. Here a weighted sum of the input variable *x*_*i*_, *i* ∈ {1, … , *M*} (reward history) is assumed to be a logit function of the dependent binary output variable *y* (return choice). To estimate the weights *β*_*i*_ for each element of the reward history, the weighted sum *h*(*x*) is computed,

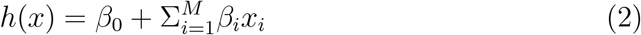

and used to define

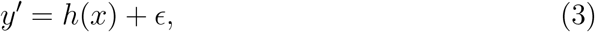

where *ϵ* is the remaining difference (error) between *y*′ and the estimate of *y*′, *h*. *y*′ is a continuous latent variable that needs to be mapped to the binary output variable *y*. Thus, the probability of seeing *y* = 1 is a logistic function of *h*,

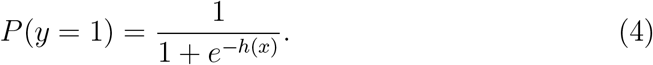

Logistic regression yields estimates of the parameters *β*_*i*_ from the data which can be used to make predictions.

To understand the values of the regression weights and what can be concluded about the fly behavior from them, we generated 100000 element reward vectors with different reward probabilities (5-30%). Under the assumption that the regression weights are determined by how often the flies returned to a stimulation and neglecting any reward correlations, we generated corresponding return choice vectors. The percentage of return choices following a reward was set to approximate ‘medium responsiveness’, with 50% correspondence. There were no choices on unrewarded trials. The regression weights can be seen in Fig. S3.

### Reinforcement Learning Models

The following reinforcement learning models [5] were applied to the data to identify potential underlying algorithms: a Rescorla Wagner (RW) model [6], a forgetting model where learning and forgetting rates are equal (termed FQ model) and a forgetting model where learning and forgetting happen at different rates (termed FQ^*αF*^).

The RL models were fit to each individual fly using maximum likelihood estimation with the following log likelihood function

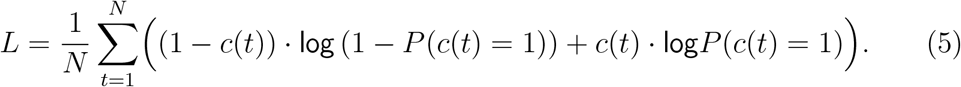

*c*(*t*) = 1 corresponds to a return choice and *c*(*t*) = 0 corresponds to no choice on trial *t*. The simple RW model has three parameters, *α*, *β* and *bias*, where *α* is the learning rate, determining the impact of the reward-prediction error, *R*(*t*) − *Q*(*t* − 1), on the value update, where *R*(*t*) is the reward at trial *t* and *Q*(*t*) is the value corresponding to a choice. *β* is the weighting factor of the value in the choice probability,

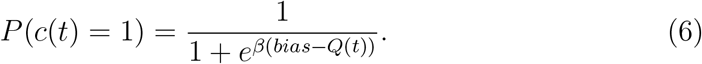

A *bias* parameter was included, to account for the fact that the baseline return probability for a fly is below 50%. In this simple model, the value of a choice *c* = 1 is only updated, when the fly made a choice, and remains constant otherwise (*ϕ* = 0 in Eq.). To make the model slightly more realistic, a second RL model, the FQ model, was implemented, where the value of a choice was forgotten, if the fly didn’t make a choice, with the same learning parameter *ϕ* = *α* as in the value update equation. The third model had one additional parameter, a forgetting parameter *ϕ* = *α*_*F*_, to allow for the more general case of different strengths of the learning and forgetting processes.

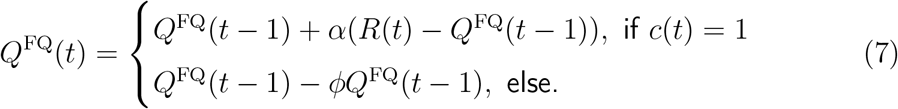

Every fly was fit with 100 random initializations of these parameter sets for each model and the best parameters were selected by the corresponding highest log likeli-hood values, ln(*L*). Subsequently, the Akaike Information Criterion [53] (AIC) score was computed, to select the one that best fits the data, while taking the number of parameters into account.

To allow for predictive testing of the models, only half of every fly data was used to fit the parameter values and the other half was used to predict the flies’ choices. The *F*_1_ score was used as accuracy measure for every fly,

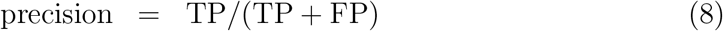

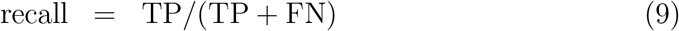

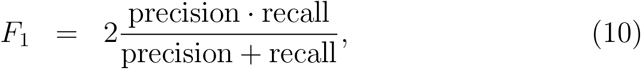

with TP the rate of true positives, FP the rate of false positives and FN the rate of false negatives.

To test the models’ generative power, 1000 sequences of 1000 trials each for the different experimental probability conditions were simulated.

## Acknowledgement

This study was supported by Lundbeckfonden grant no. DANDRITE-R248-2016-2518. We gratefully acknowledge helpful comments from Dennis Eckmeier, Alex Gomez-Marin, Daisuke Hattori and Ollie Hulme.

## Author Contributions

JIS and DK designed the experiment. SES performed all experiments and analyses. SES and DK wrote the manuscript. ACvP provided expertise and feedback.

## Supplementary Information

### Supplementary Figure 1

The optogenetic stimulation changes the walking patterns of the flies, from a more uniform positional coverage of the arena, to a stimulation zone localized occupancy (Fig. S1A,B). The speed distribution of genetic and unstimulated control flies is bimodal, with a slow peak from wall approach and a fast peak from walking in the inner part of the arena, which is also (but to a lesser degree) preserved on the unrewarded trials of the stimulated populations (Fig. S1A left). Upon stimulation, the fast peak is decreased and the slow peak slightly increased (Fig. S1A. right). The average walking speed is similar across conditions and strongly reduced in contrast to freely walking flies [1], due to the confinement (Fig. S1D). To characterize the trials we imposed on the data, we looked at the trial length distributions of stimulated and unstimulated trials. Fig. S1C shows 4 example conditions and their population trial length distributions. Stimulated or rewarded trials were longer due to the lingering time from experiencing the reward and the subsequent local search like behavior. The distributions are similar across probability conditions, indicating that the arena geometry and thereby the walking speed are imposing boundaries on the typical duration of a trial. Since the population speed distributions in Fig. S1A didn’t show a very clear effect of the stimulation on the walking speed, we looked at the local walking speed distribution of the flies after they received a reward. When walking out of the stimulation zone, the flies showed a fast peak, while when they then returned to the stimulation zone, the speed was reduced (Fig. S1D). This effect was averaged out in the population speed distribution.

**Figure S1:**
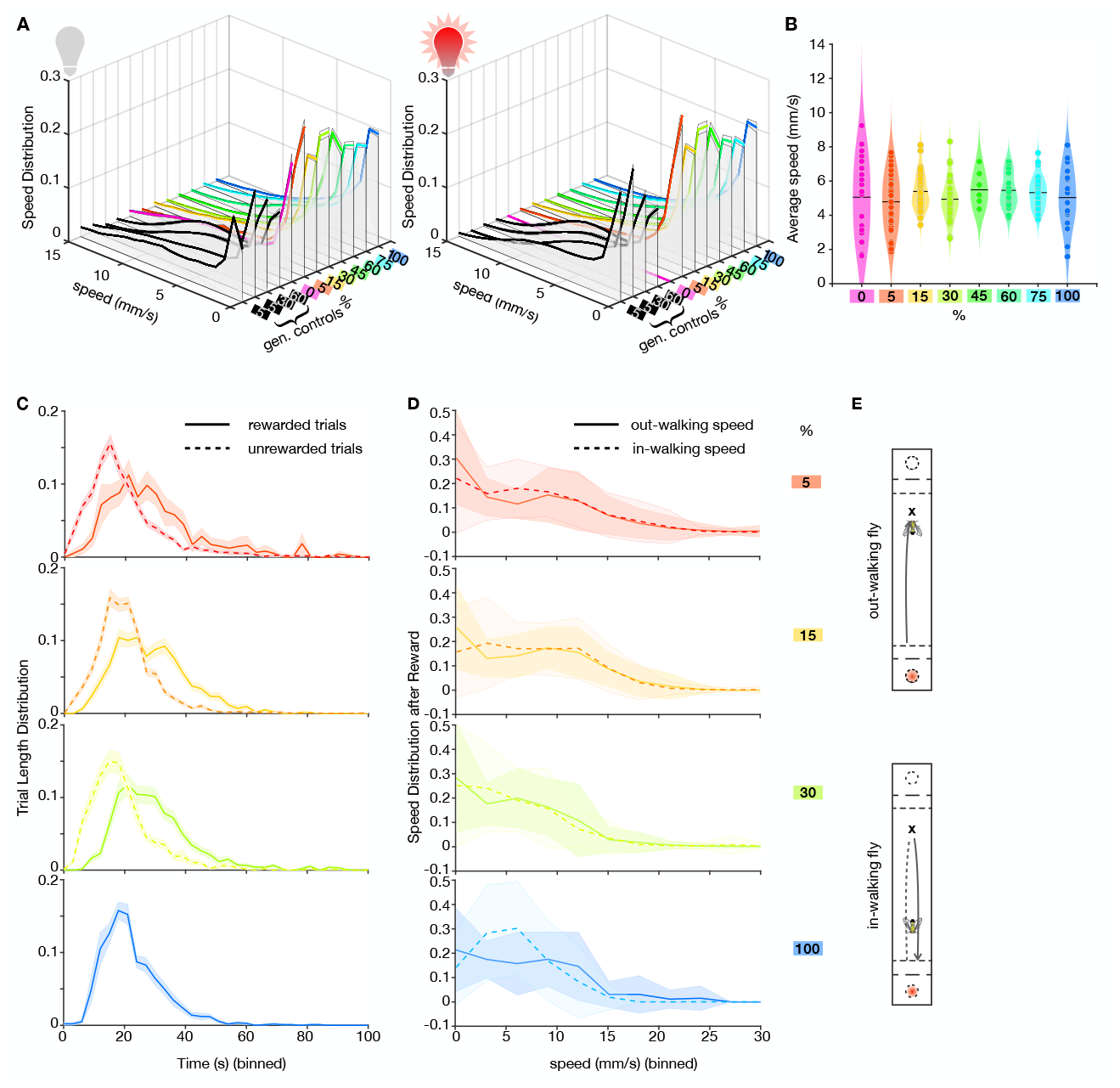
Population speed and Characterization of Trials. **A** Speed distribution of fly populations. Left: unstimulated trials (grey light bulb). Right: stimulated trials (red light bulb). Stimulation affects the higher speeds but not the low speeds. **B** Average walking speed per condition. **C** Trial length distribution of 5,15,30 and 100% condition populations. Solid lines show rewarded trials (longer) and dashed lines show unrewarded trials (shorter). **D** Speed distribution after a reward. Solid lines: out of the reward zone walking speed. Dashed lines: in-walking speed when returning from walking out (same trial as out-walking speed). In-walking is on average slower than out-walking. **E** Pictogram of out-walking and in-walking traces.

### Supplementary Figure 2

In addition to the single sided experiments, we also collected double-sided stimulation data in two conditions, 5:5% and 15:15%. We performed the same analysis on this data as on the single-sided data. The occupancy distribution showed peaks in both stimulated zones (Fig. S2A) and the zone preference indices were close to zero in both cases, indicating no preference for one zone over the other (Fig. S2B). The returns to both zones were not significantly different and similar to those of the single-sided conditions of the same probability (Fig. S2C). Since the first reward could happen in either zone, we compared first reward return probabilities to both zones for the two double-sided conditions (Fig. S2D left) and the corresponding first rewarded trial numbers (Fig. S2D right). Both were consistent with the results from the single-sided cases. Logistic regression analysis (Fig. S2E) revealed again that the current reward was most predictive of a return and that there was no influence of the return history. To test whether the animals also followed a reinforcement learning algorithm to allocate their choices to two options, we used the same three types of models, extended by a second option. Model comparison (Fig. S2H) yielded the lowest AIC score for the FQ model, which captured return probabilities to both zones, as well as return run lengths in both probability conditions in a generative test, well (Fig. S2J,K).

**Figure S2:**
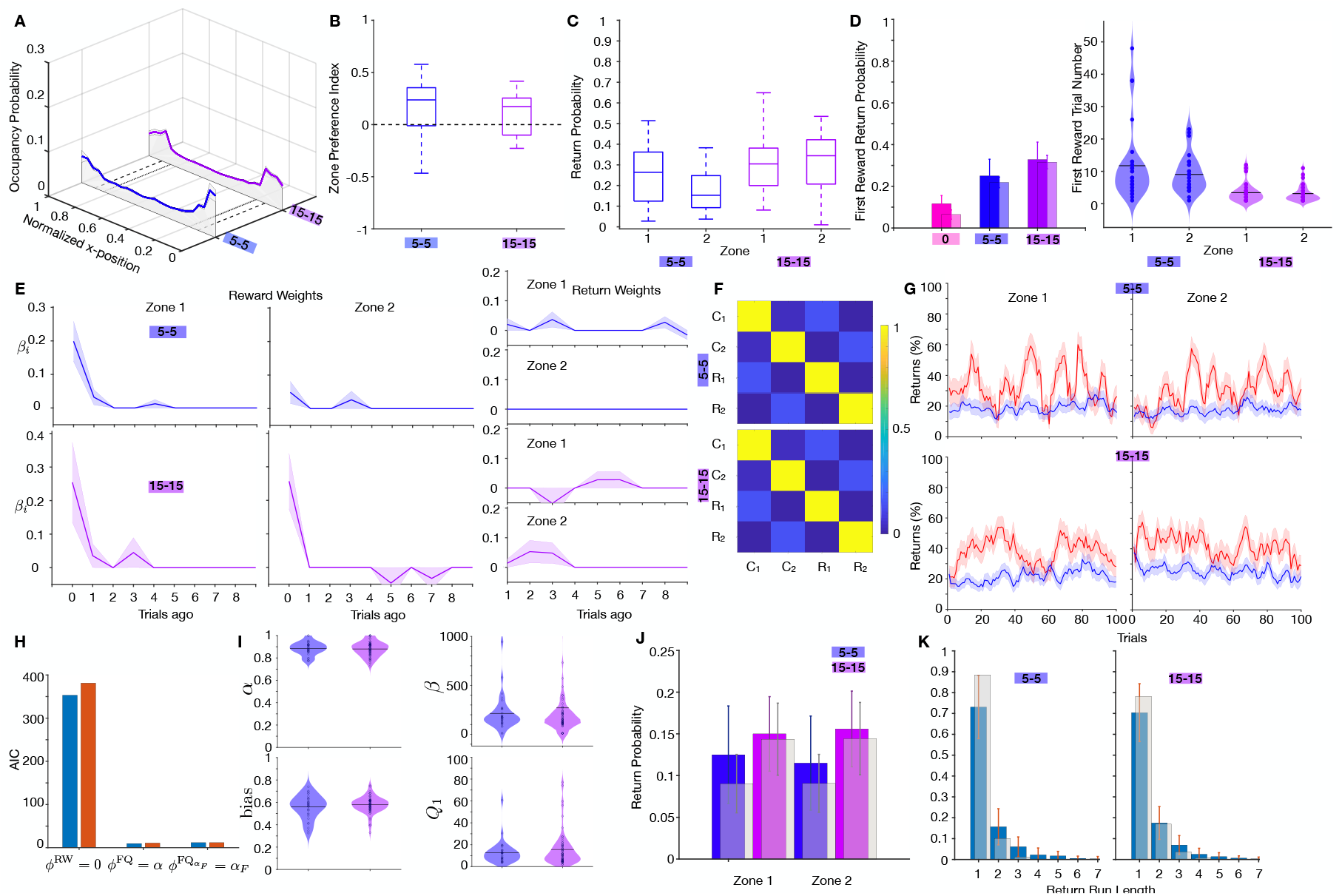
Experiments with double-sided stimulation. **A** Occupancy distribution for 5% and 15% double-sided stimulation data. **B** Preference index. **C** Return probability to zones 1 and 2 for both conditions. **D** Left: Return behavior on the first reward (to either zone) compared to the first trial return for unstimulated controls. Right: First rewarded trial number. **E** Left and center: Logistic regression weights of returns against the reward history for both population data to each zone independently. Right: Logistic regression weights for returns against return choice history. **F** Pearson correlation for rewards (R) and returns (C). **G** Return behavior as 5-trial moving average. Red curves: rewarded trials, blue curves: unrewarded trials to the same zone. **H** AIC score for the three RL models. **I** Best-fit parameter values of the FQ model. **J** Generative testing of the FQ model: comparison of the return probability (exp. data: colored, model: grey). **K** Generative testing of the FQ model: Return run lengths (exp. data: blue, model: grey).

### Supplementary Figure 3

We concluded that the flies perform local searches after receiving a reward since their turns increase around the reward location. How do those distributions look in time after a reward? Fig. S3A shows population averaged turns for individual rewarded (upper row) and unrewarded (bottom row) trials in time since a trial start at t = 0. Each local search time was cutoff at the time of the return to the reward zone. To capture the general time course, both scatter plots are summarized as histograms in the middle row. Comparing those distributions shows that rewarded flies perform more temporally extended curved trajectories than unstimulated flies. We performed the same analysis on the time points of returns, Fig. S3B and revealed that, if they return, unstimulated trials are returned faster than rewarded trials, consistent with a more extended local search behavior after a reward. Are those local searches a hard-wired behavior that is always elicited upon reward encounter (specifically sweet taste rewards) or do they undergo adaptation to the reward probability? To test this, we show polar plots of the angular distribution of the walking paths in the reset zone in Fig. S3C. Rewarded (solid lines) and unrewarded (dashed lines) trials separate in this visualization, since the search path has a larger variability in turn angles than an unrewarded fly’s path, that only turns at the arena wall. While they are significantly different within the probability condition, the angular distributions on rewarded trials across probability conditions are not. Furthermore, the unrewarded angular distributions across conditions are significantly different from the unstimulated control flies (0%). Together, this suggests that local searches emerge upon reward encounter, they are hard-wired in the sense that they are not different across probability conditions and unrewarded trials actually have less ‘curvy’ paths than those of always unstimulated flies. Providing a second reward location, as in the double-sided conditions, can help elucidate whether the flies localize their returns in space to the availability of rewards in zone 2 of the arena. We compared the distribution of returns in space from the reward zone for the 5 and 15% single and double-sided data, Fig. S3D. In all conditions, the flies are more likely to return spatially close to the previously visited reward zone and the probability decreases the further away the fly walks. There was no difference between the single- and double-sided conditions, rejecting our hypothesis of return localization.

**Figure S3:**
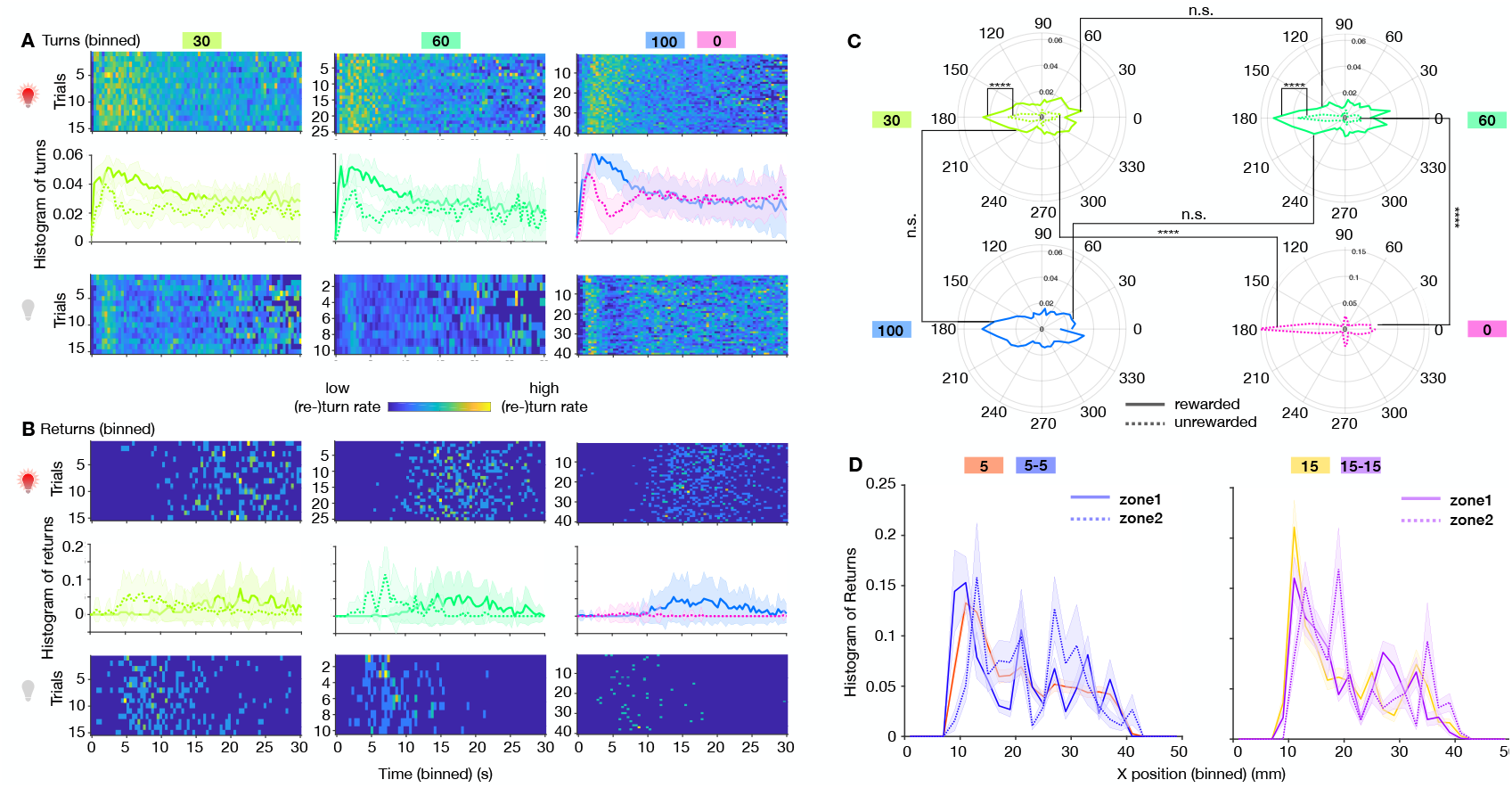
Local search analysis. **A** Top row: Turns, as a proxy for local search, (binned) in time since trial start for 30, 60 and 100/0% conditions. Middle row: histogram of temporal turn distribution. Solid drawn curve corresponds to rewarded trials (top row) and dashed curve corresponds to unstimulated trials (bottom row). Bottom row: turns in time since trial start for unstimulated trials. **B** Top row: returns on rewarded trials in time since trial start for the same fly populations. Middle row: histogram of returns. Solid curve: stimulated returns, dashed curve: unstimulated returns. Bottom row: Unstimulated returns. **C** Polar plots of angular distributionns.s. n.s. of walking traces in the reset zone, for 30%, 60%, 100% populations and unstimulated controls (clockwise). Solid lines: rewarded trials, dashed lines: unrewarded trials. (* * **: p < 0.0001, two-way Kolmogorov-Smirnoff test.) **D** Comparison of return location (maximum position of a trial) for 5 and 15% single and double sided condition fly populations. Double sided cases have rewards in both zones and thus returns to both zones are separated.

### Supplementary Figure 4

To determine how stable the return behavior was over the time of the session, we looked at 5-trial moving averages of the returns for the fly populations in each probability condition (Fig. S4A). With increasing stimulation probability, the returns upon rewards (red curves in Fig. S4A) decreased over time (trials). The data shown corresponds to 2 hours of experiment. We therefore reduced the data to 30 minutes where the return behavior was approximately stable for all conditions. To justify the regression analysis we looked at the pearson correlation of the rewards and the animals’ returns (Fig. S4B). To help interpret the logistic regression weights, we simulated rewards with 5 different reward probabilities (5-30%) and return vectors, where the return probability upon a reward was set to 50% (Fig. S4C). The size of the first coefficient was thus determined by the stimulation probability and explains the effect we see in Fig. 2D.

**Figure S4:**
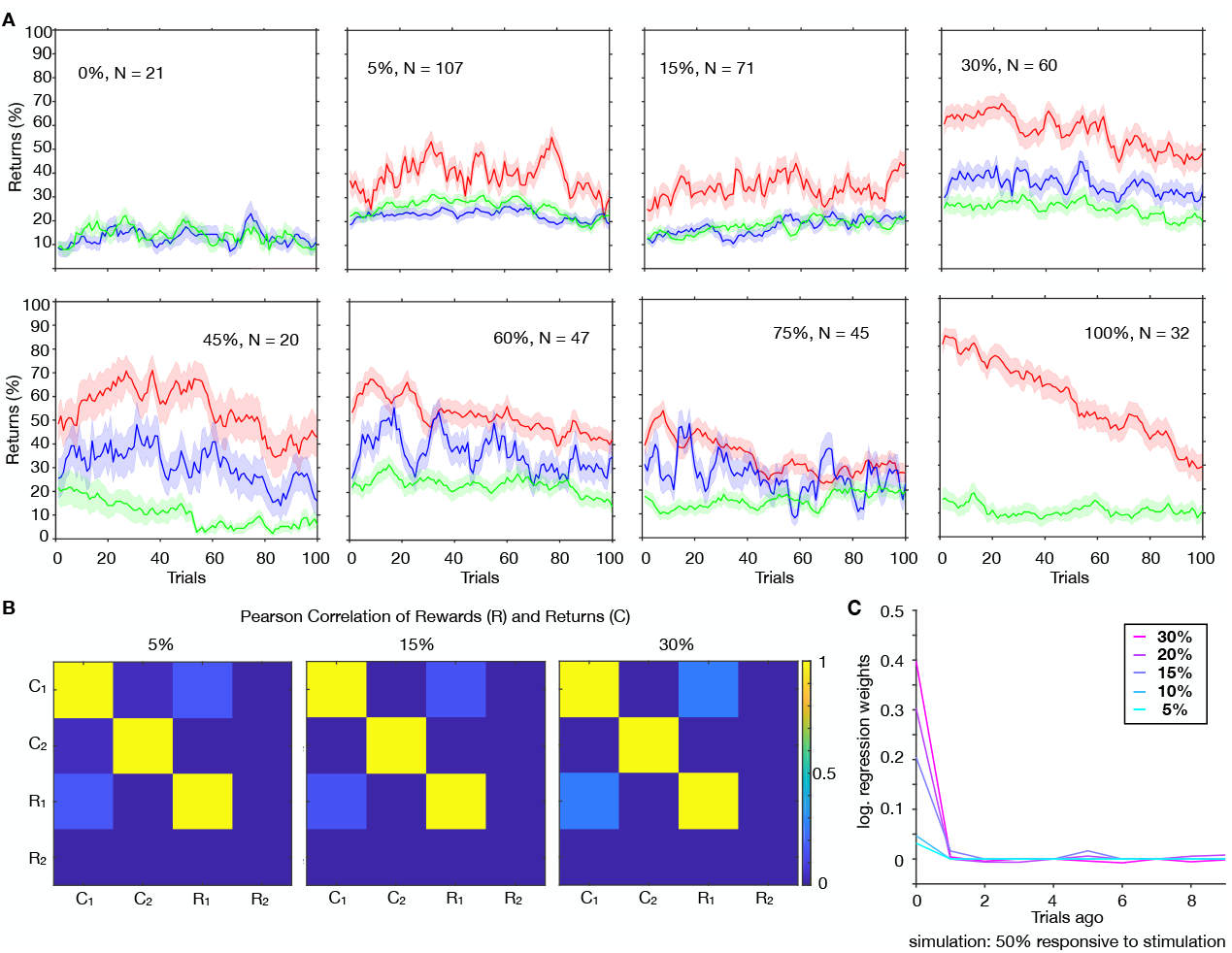
Stability of return behavior over trials and reward-choice correlations. **A** 5-trial moving averages of the returns over trials for 0-100% stimulation probability conditions. Red curves show returns upon rewards, blue curves show returns to the rewarded zone without rewards and green curves show returns to the unstimulated zone. **B** Pearson correlation of rewards and returns (choices) for 5,15 and 30% conditions. **C** Logistic regression of simulated data to rewards. Simulated data was generated with 50% return probability upon a reward. Curves show regression weights for different stimulation probabilities (5-30%).

### Supplementary Figure 5

We tested three reinforcement learning models (see also Sec.) and compared their AIC scores as a measure of how well they captured the data (Fig. S5A). The models were fit on half of the data (of each fly) and the other half was used to perform a predictive test (Fig. S5C). Especially the low probability data could not be very well predicted, which is due to the limited number of reward and return events in the data. This is visualized in Fig. S5D by means of the *F*_1_ score, a measure of predictive accuracy against the probability of returned trials. Data with more returns could be fit more reliably and yielded a higher *F*_1_ accuracy. The slightly better predictive performance of the FQ model than the RL model made us choose this model to explain the return behavior. The corresponding best-fit parameters of the flies whose behavior could be predicted by the FQ model, are shown in Fig. S5B. We furthermore used the models for generative testing, where we used the return probability and the return run length distributions as measures for comparison with the experimental data. All models perform well and generate distributions quite close to that of the exp. data.

**Figure S5:**
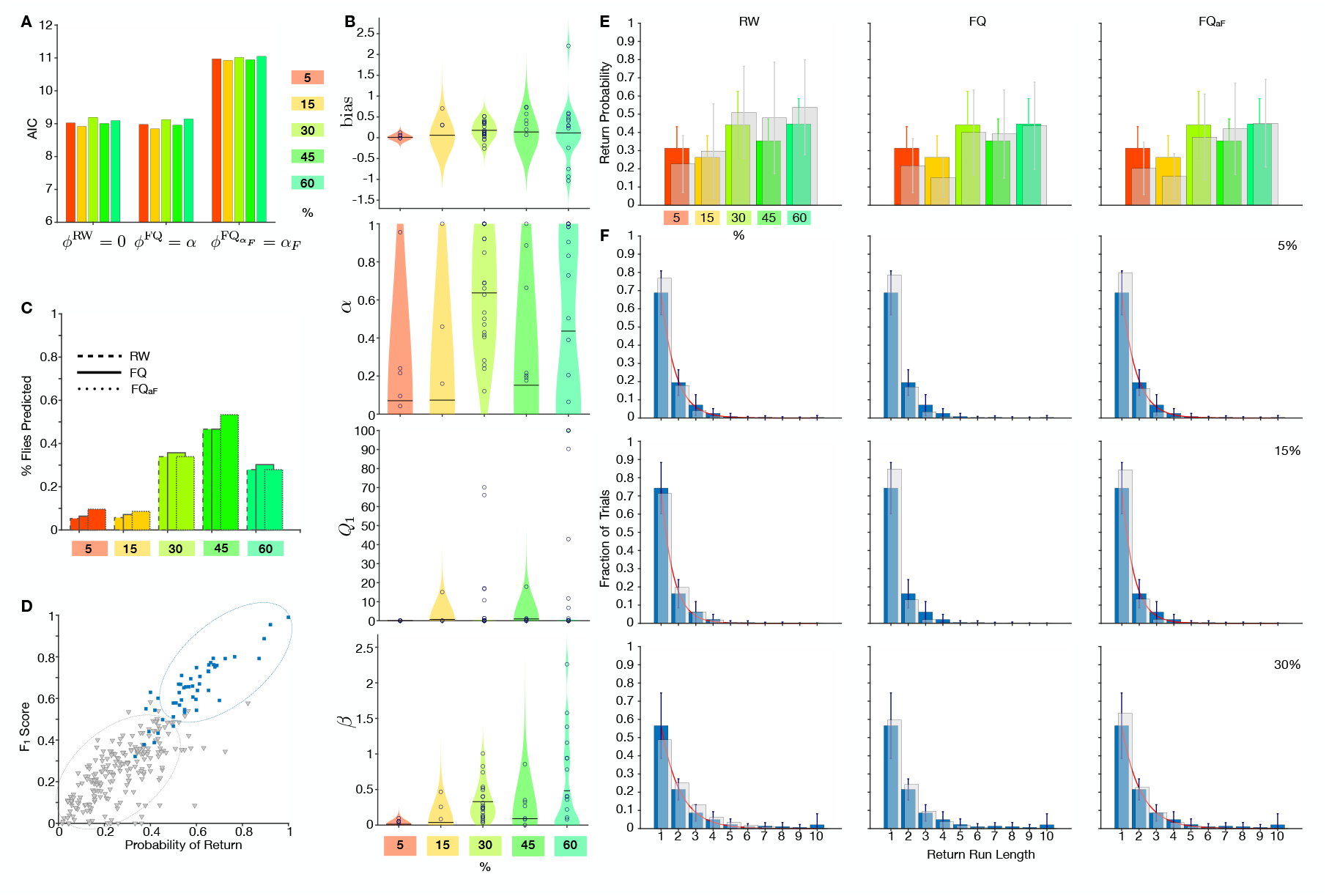
Reinforcement learning model selection, predictive test and generative test. **A** AIC scores for the three RL models on 5-60% data. The lower the AIC score, the better the model captures the data while excessive parameters are punished. **B** Best-fit parameter values of the FQ model for each fly (circles) and population averages (solid lines in the violins). **C** Predictive test of the FQ model. Number of flies that could be predicted with more than 50% accuracy (*F*_1_ score) for each model. Total number of flies per condition: *N*^5%^ = 94, *N* ^15%^ = 70, *N* ^30%^ = 56, *N* ^45%^ = 15, *N* ^60%^ = 45. **D** *F*_1_ score against data choice probability. If choices made up less than 50% of the data, the model had a poor predictive power. Dashed ellipses visualize clustering of the data with high and low *F*_1_ score. **E** Comparison of generative properties of the three RL models: Return probability. **F** Comparison of generative properties of the three RL models: Return run lengths. Red curves: exponential fits.

